# Prenatal THC does not affect female mesolimbic dopaminergic system in preadolescent rats

**DOI:** 10.1101/2020.12.23.424157

**Authors:** Francesco Traccis, Valeria Serra, Claudia Sagheddu, Mauro Congiu, Pierluigi Saba, Gabriele Giua, Paola Devoto, Roberto Frau, Joseph Francois Cheer, Miriam Melis

## Abstract

Cannabis use among pregnant women is increasing worldwide along with permissive sociocultural attitudes towards it. Prenatal cannabis exposure (PCE), however, is associated with adverse outcome among offspring ranging from reduced birth weight to child psychopathology. We have previously shown that male rat offspring prenatally exposed to Δ9-tetrahydrocannabinol (THC), a rat model of PCE, exhibit extensive molecular, cellular and synaptic changes in dopamine neurons of the ventral tegmental area (VTA), resulting in a susceptible mesolimbic dopamine system associated with a psychotic-like endophenotype. This phenotype only reveals itself upon a single exposure to THC in males but not females. Here, we characterized the impact of PCE on female behaviors and mesolimbic dopamine system function by combining in vivo single-unit extracellular recordings in anesthetized animals and ex vivo patch clamp recordings, along with neurochemical and behavioral analyses. We find that PCE female offspring do not show any spontaneous or THC-induced behavioral disease-relevant phenotypes. The THC-induced increase of dopamine levels in nucleus accumbens was reduced in PCE female offspring, even when VTA dopamine activity in vivo and ex vivo did not differ compared to control. These findings indicate that PCE impacts mesolimbic dopamine function and its related-behavioral domains in a sex-dependent manner and warrant further investigations to decipher the mechanisms determining this sex-related protective effect from intrauterine THC exposure.

**Highlights:**

- PCE female offspring do not manifest a disease-relevant phenotype
- Prenatal THC does not affect female dopaminergic neurons
- PCE female mesolimbic dopamine function is less responsive to acute THC

## 1. Introduction

The prevalence of mental disorders worldwide is 13% and, alarmingly, 50% of these are established before the age of 14 years with most cases undetected and untreated ^1,2^. Additionally, the average delay between symptom onset and initial treatment is of about 11 years ^3^, a delay that potentially jeopardizes clinical outcome. This highlights the relevance of an early identification of these disorders to design a timely therapeutic intervention.

Children with a parent who has a mental illness or substance use disorder have a higher risk of psychiatric problems themselves ^4,5^. Critically, prenatal cannabis exposure (PCE) increases the risk for child psychopathology, ranging from affective symptoms to ADHD and psychotic-like experiences ^6–11^. With recreational cannabis legalization and permissive sociocultural attitudes expanding worldwide, and the use of cannabis among pregnant women on a sharp rise ^12–14^, concern increases over the long-term negative impact on next generation health (i.e., pediatric concern)^15–19^. Since cannabis main psychoactive ingredient Δ^9^-tetrahydrocannabinol (THC) crosses the placenta and interferes with the endocannabinoid system, a signaling pathway key in proper neural development ^20–23^, it is plausible that PCE may be teratogenic ^12^. Indeed, PCE is thought to act as a “first hit” on neurodevelopmental trajectories ^24^. Of note, a sex bias in response to PCE is well recognized ^6–8^. However, the underlying mechanisms are not sufficiently studied also because causal inference is difficult to be established in population studies. Additionally, gender as a variable in the susceptibility to the consequences of cannabis exposure on neurocognitive and behavioral development of the offspring is seldom examined ^25^. Thus far, insights into the mechanisms of how sex interacts with PCE to generate unique effects in the progeny can be solely gained from preclinical studies.

Animal models of PCE show that males exposed offspring are more susceptible than females to dysfunctions in cognitive processing and emotional regulation ^26–35^. Thus, female sex often is a protective factor in response to same intrauterine environmental insults. However, many individual (e.g., species, strain, age) and experimental (e.g., design, drug, dosage, route, regimen) variables along with objective endpoints (e.g., behavioral paradigm, experimental technique) might influence sex dimorphism. For instance, we have shown that only male PCE offspring exhibits, at pre-puberty, a psychotic-like endophenotype^34^. This is accompanied by extensive molecular, and synaptic changes in dopaminergic neurons of ventral tegmental area (VTA), converging on a mesolimbic hyperdopaminergic state susceptible to either THC or stress ^34,36^. However, whether female offspring display a different disease-relevant behavioral phenotype likely associated with yet unidentified alterations in mesolimbic dopamine system function remains unknown. Here we show that many of the detrimental effects induced by PCE on male dopamine neurons and their neurochemical and behavioral readout are absent in females at prepuberty. Of note, PCE female progeny manifest a risk-safer phenotype along with normal social behavior and adaptive coping strategies to acute stress. Collectively, this data extends our understanding of the multifaceted developmental effects imposed by PCE on the rat mesolimbic dopamine system and warrant further investigations to decipher the mechanisms determining this process that confers a sex-specific resistance against prenatal THC.

## 2. Results

### 2.1 Impact of PCE on behavior in female preadolescent rats

Evidence suggests a sex bias in response to PCE ^6–8^. In particular, we found that male PCE rats display a psychotic-like phenotype in response to acute THC or stress, a higher sensitivity to a dopamine D2 agonist and engage in risk taking behaviors ^34,37^. However, female progeny does not exhibit sensorimotor gating deficit in response to PCE or acute THC ^34^. Therefore, we first evaluated whether female progeny displayed other features of PCE male cohort.

When we measured spontaneous or THC-induced locomotor activity in vehicle (CTRL) and PCE female offspring in an open field arena, no differences were observed at baseline (two-way ANOVA, main effect of PCE: F_(1,30)_= 1.67, P=0.206) and following an acute administration of THC (2.5 mg/Kg, s.c.; Figure 1a; two-way ANOVA, interaction, F_(1,30)_= 3,336, P=0.1; main effect of THC: F_(1,30)_= 5.384, p=0.027). Accordingly, no differences were observed in thigmotaxis between groups (Figure 1b; two-way ANOVA, interaction PCExTHC: F_(1,29)_= 0.091, p=0.7645). Next, we investigated risk propensity by using the wire-beam bridge task, which evaluates a risk-taking phenotype in rodents by measuring their proclivity to cross a flexible bridge suspended over a 150 cm deep gap. PCE increased the latency to cross the wire-beam bridge (Figure 1c; two-way ANOVA, main effect of PCE: F_(1,26)_= 6.306, p=0.018), but did not affect the response to acute THC (main effect of THC: F_(1,26)_= 0.031; p=0.859) or the number of stretched-attend postures (Figure 1d; two-way ANOVA, main effect of THC: F_(1,26)_=4.987, p=0.034; interaction PCE x THC: F_(1,26)_=0.091, p=0.76). Since the spontaneous prudent phenotype observed in PCE females could be secondary to an anxiety-like state, we carried out the elevated plus maze test. In agreement with stretched-attend posture phenotypes (as index of anxiety-like phenotype) observed in the bridge test, we found no differences between the groups (Figure 1e; two-way ANOVA, interaction: F_(2, 41)_=0.776, p=0.467). Finally, given that PCE male progeny do not adopt copying strategies in response to acute stressors ^37^, we evaluated the effects of forced swim test, as an acute inescapable stressor, on female offspring. PCE does not alter the amount of time spent in engaging active (struggling/climbing and swimming) or passive (immobility) strategies (Figure 1f; two-way ANOVA, interaction: F_(2, 48)_=0.293; p=0.7475). Collectively, these results indicate that PCE does not induce in female offspring a susceptible phenotype resembling their male counterparts.

**Figure 1.**
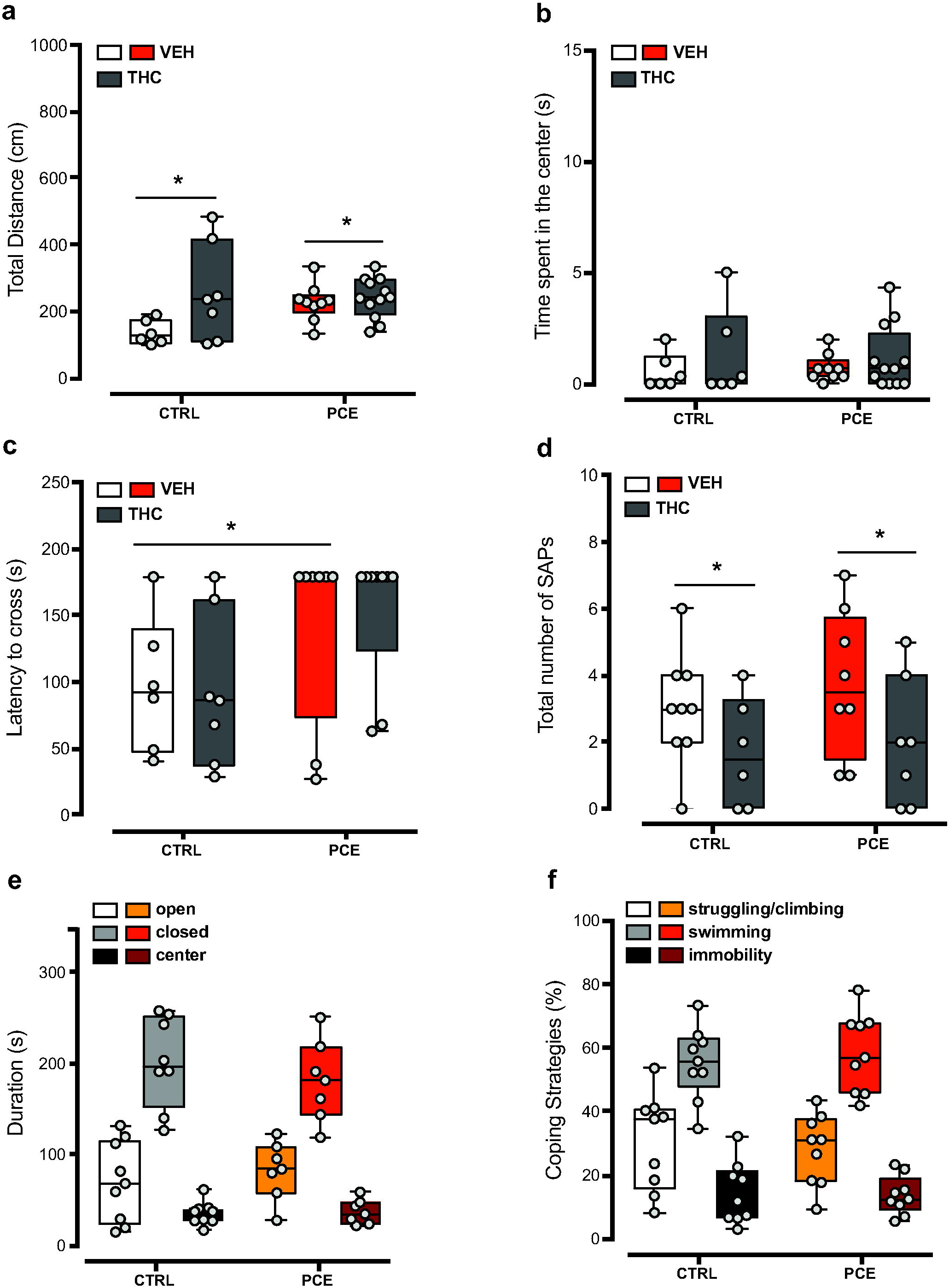
PCE effect on female preadolescent offspring behavior. (**a**) Spontaneous locomotor activity in female offspring, measured as total distance travelled in a novel open field arena. Dose of THC is 2.5 mg ^−1^ Kg (s.c.). (n_rats_=6-12/group) (**b**) Time spent (s) in the center part of the open field arena. THC or VEH administration do not affect the thigmotaxic behavior of PCE females in comparison to CTRL. (**c**) The latency to crossing the wire-beam bridge is increased by PCE (n_rats_=7-9). (**d**) THC-challenge decreases the number of stretched-attend postures (SAP) in both PCE and CTRL offspring during the wire-beam bridge test. (**e**) Behavioral responses in the elevated plus maze. No differences were found in the total duration of time spent by offspring in the closed, open arms and center position (n_rats_= 7-9/group). (**f**) Effect of forced swim tests (FST) on coping strategies engaged by PCE and CTRL offspring (n_rats_=9/group). Time spent (%) in struggling/climbing, active swimming and immobility during the FST. All data are represented as box and whisker plot with single values (min to max). * p < 0.05, ** p < 0.01.

We next considered the possibility that PCE might induce different disease-relevant phenotypes in female progeny. Thus, we investigated whether PCE female offspring might exhibit three traits of maladaptive behaviors that occur in several psychiatric disorders, the avoidance of aversive or threatening stimulus, the social behaviors and the ability to fell pleasure towards positive emotional states. We measured the passive avoidance latency during the training session and 24 h later (i.e., retention): although an effect was observed during the training session (Figure 2a; two-way ANOVA, Bonferroni’s multiple comparisons test CTRL vs PCE, p > 0.0064), no differences were found during the retention session (Figure 2a; two-way ANOVA, Bonferroni’s multiple comparisons test, CTRL vs PCE, p > 0.999;). We then assessed whether PCE affected their social capabilities, but found no differences between the groups, in terms of duration of both nonsocial and social activities during social interaction test (Figure 2b; two-way ANOVA, interaction: F_(2, 42)_=1.789, p=0.179). Specifically, PCE did not alter the frequency and the duration of social exploration, measured as sniffing approaches to the partner (Figure 2b-**c**; frequency: unpaired t-test, t_15_=1.322, p=0.205; duration: t_15_=1.52, p=0.149). Accordingly, no differences were found in the frequency of pinning and pouncing (Figure 2c; unpaired t-test, pinning: t_14_=0.355, p=0.72; pouncing: t_14_=0.550, p=0.59) and in total time spent in playing behaviors (Figure 2d; t_14_=0.606, p=0.55).

**Figure 2.**
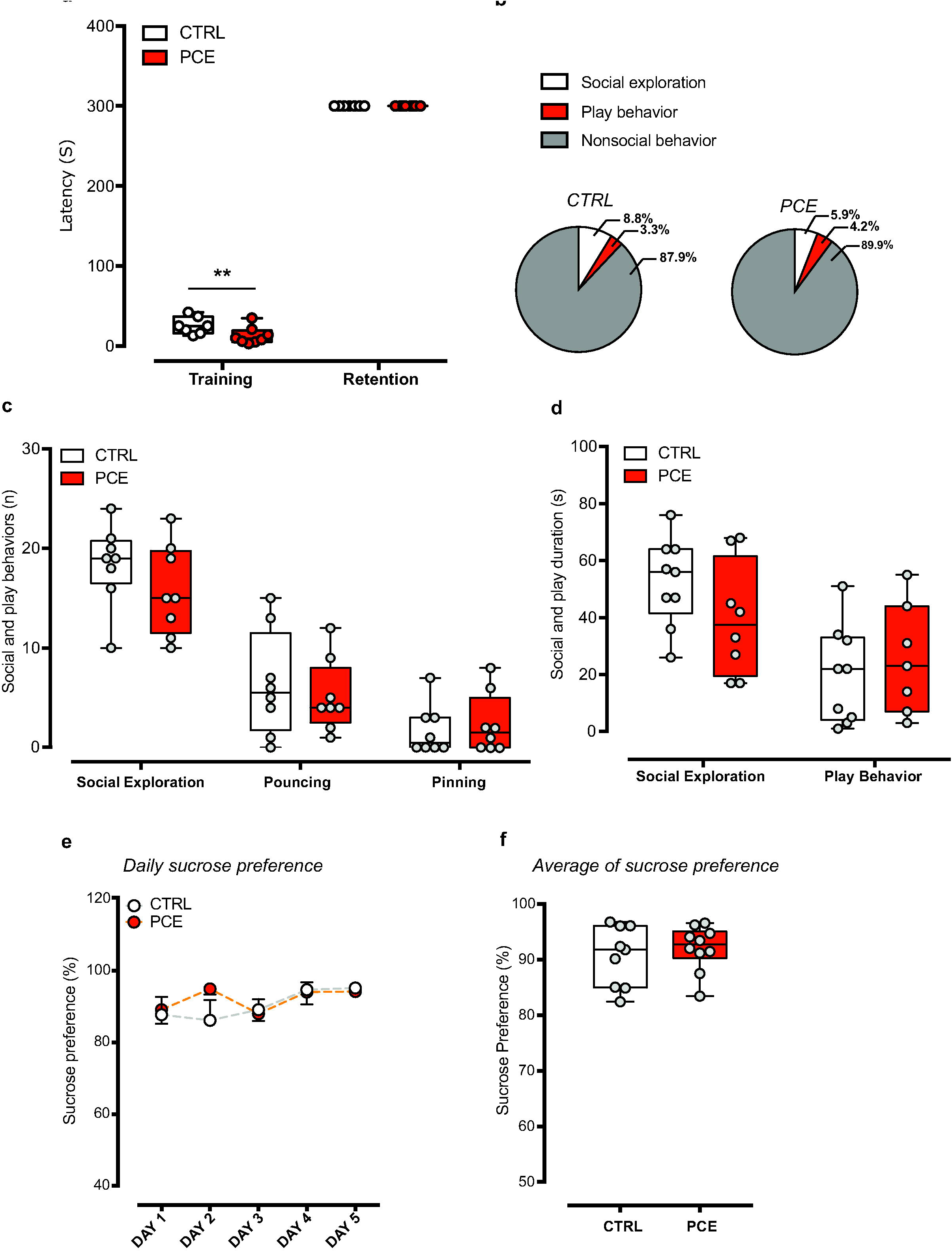
PCE impact on emotional memory, social interaction and anhedonia-like behavior in female preadolescent rats. (**a**) Passive avoidance (PA) latency (s) to enter in dark compartment during training (before foot shock) and retention (24 h after foot shock) sessions (n_rats_=7-8/group). (**b**) Pie chart representation of social (exploration and play behavior) and nonsocial behaviors exhibited during the social interaction test. No differences were found in terms of (**c**) number of social approaches, pouching and pinning and (**d**) total duration of time spent in social exploration and play behaviors (n_rats_=7-9/group). (**e**) Daily sucrose preference (%) during the 5 days of sucrose preference testing and (**f**) in the average sucrose preference. Data are represented as average ± s.e.m (for sucrose preference curves) or as box-and-whiskers plot with single values (min to max). **, p < 0.01.

Finally, we investigated whether PCE affected the animals’ ability to experience pleasure by performing the sucrose preference test (SPT), whose changes are suggestive of anhedonia, a cardinal symptom in depressive states. PCE did not affect female offspring daily sucrose preference (Figure 2e; RM two-way ANOVA, F_(4, 83)_ = 0.863, p=0.4893) either at baseline (day 1) and during the subsequent days (from day 2 to 5). Accordingly, no differences were found in the average preference for sucrose solution (Figure 2f; unpaired t-test, t_17_=0.664, p=0.51). Altogether, PCE does not impact any of the behaviors studied here in females.

### 2.2 PCE effect on mesolimbic dopamine transmission

To elucidate whether the apparent protection from PCE impact observed in female progeny may be related to sex differences in the effects of PCE on mesolimbic dopamine system function, we next carried out cerebral microdialysis experiments in the target region of the shell of nucleus accumbens (NAcS) (Figure 3a,b). In behaving pre-adolescent female offspring, we found no alteration in basal extracellular dopamine levels between CTRL and PCE females (Figure 2c; unpaired t-test, t_14_=0.72, p=0.48). Notably, in PCE offspring THC-induced increase in extracellular dopamine was smaller when compared to CTRL (Figure 3d; RM two-way ANOVA, PCE: F_(1, 13)_ = 5.11, p=0.04). Next, we investigated whether PCE affects electrophysiological properties of VTA dopamine neurons (Figure 4a) putatively projecting to NAcS by performing single unit extracellular recordings *in vivo*. In anesthetized rats, neither the number of spontaneously active cells (Figure 4b; unpaired t-test, t_9_ =0.05; CTRL: 1.16 ± 0.11, n=5; PCE: 1.15 ± 0.27, n=6) nor the average firing frequency (Figure 4c,d; unpaired t-test, t_85_ =0.71; CTRL: 2.71 ± 0.27, n=35; PCE: 2.96 ± 0.21, n=52) differ in PCE (n_rats_=6, n_cells_=52) as compared with CTRL (n_rats_=5, n_cells_=35). Similarly, the firing mode, as expressed by the percentage of spikes in bursts (Figure 4e; unpaired t-test, t_85_ =0.45) and pattern of activity (Figure 4f; PCE regular, n_cells_=21; irregular, n_cells_=20; bursting, n_cells_=11; CTRL regular, n_cells_=12; irregular, n_cells_=15; bursting, n_cells_=8) did not change between the groups (Chi-square test=0.33). We next examined whether dopamine neurons responded differently to the effect of an acute THC challenge alike males ^37^. We found that THC (0.5 mg/kg i.v.) did not alter the firing frequency of VTA dopamine neurons in either group of preadolescent females (n_cells_=5 in both groups; Figure 4g; RM two-way ANOVA, F_(5,40)_=0.28).

**Figure 3.**
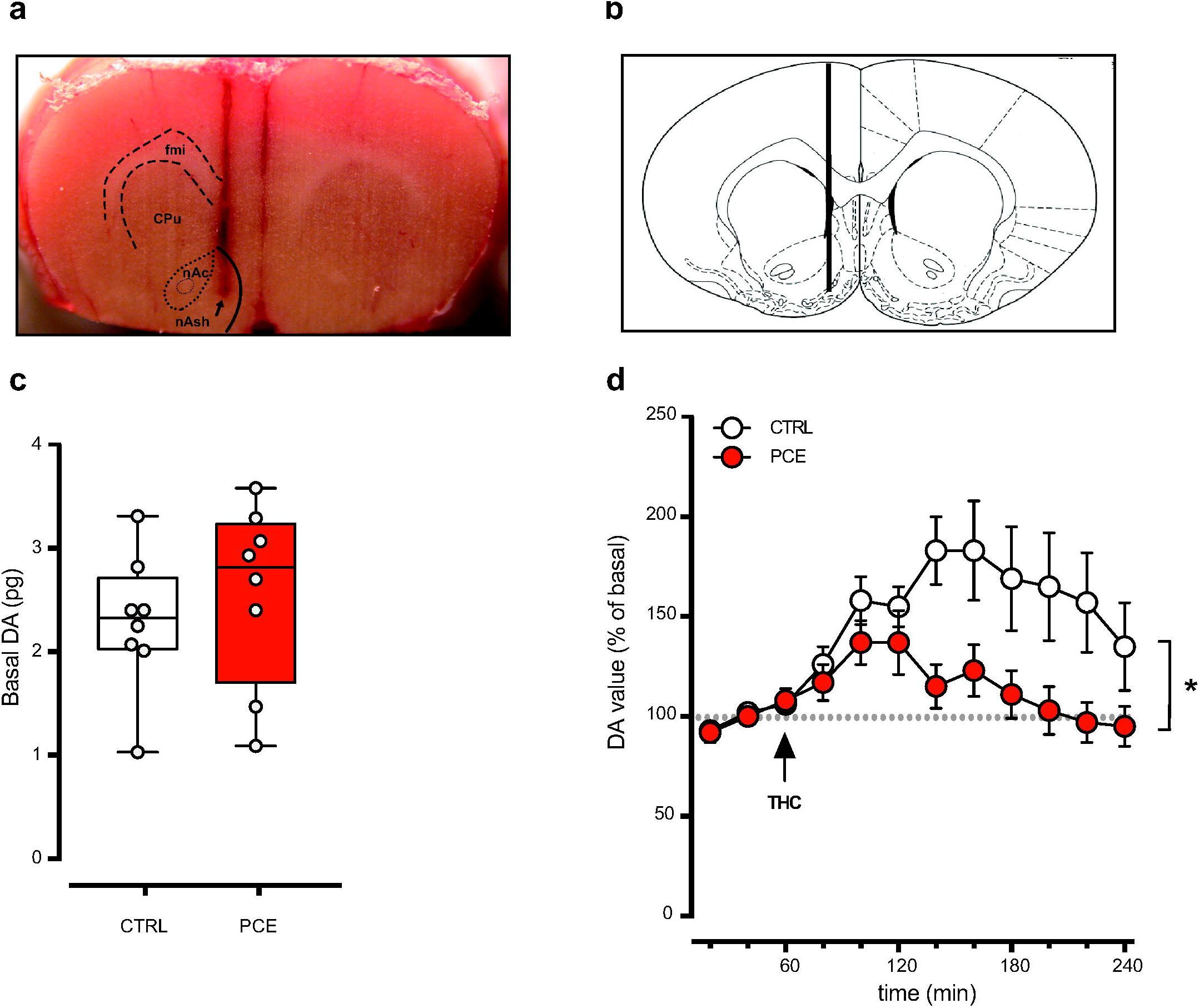
PCE impact on THC-induced increase of accumbal DA level in female preadolescent offspring. (**a**) Representative photograph of microdialysis probe location into the nucleus accumbens shell. The arrow indicates the tip of the probe. Abbreviations: fmi, forceps minor corpus callosum; CPu, caudate putamen; nAc, nucleus Accumbens core; nAsh, nucleus Accumbens shell. (**b**) Schematic representation of cerebral area targeted by the probe as indicated by the vertical line (nAsh, AP: +1.5, L: ±0.7, V: −7.0 from bregma) from the atlas ^74^. (**c**) Basal dopamine (DA) values (pg) measured in the nAsh (n_rat_=7-8/group). Data are expressed as pg/sample (box and whisker plot min to max with single values). (**d**) Time course of the effect of acute THC (2.5 mg kg-1 i.p.) on extracellular DA levels in nAsh (n_rat_=7-8/group). Data are shown as percent of baseline and are represented as mean ± s.e.m. *, p<0.05.

**Figure 4.**
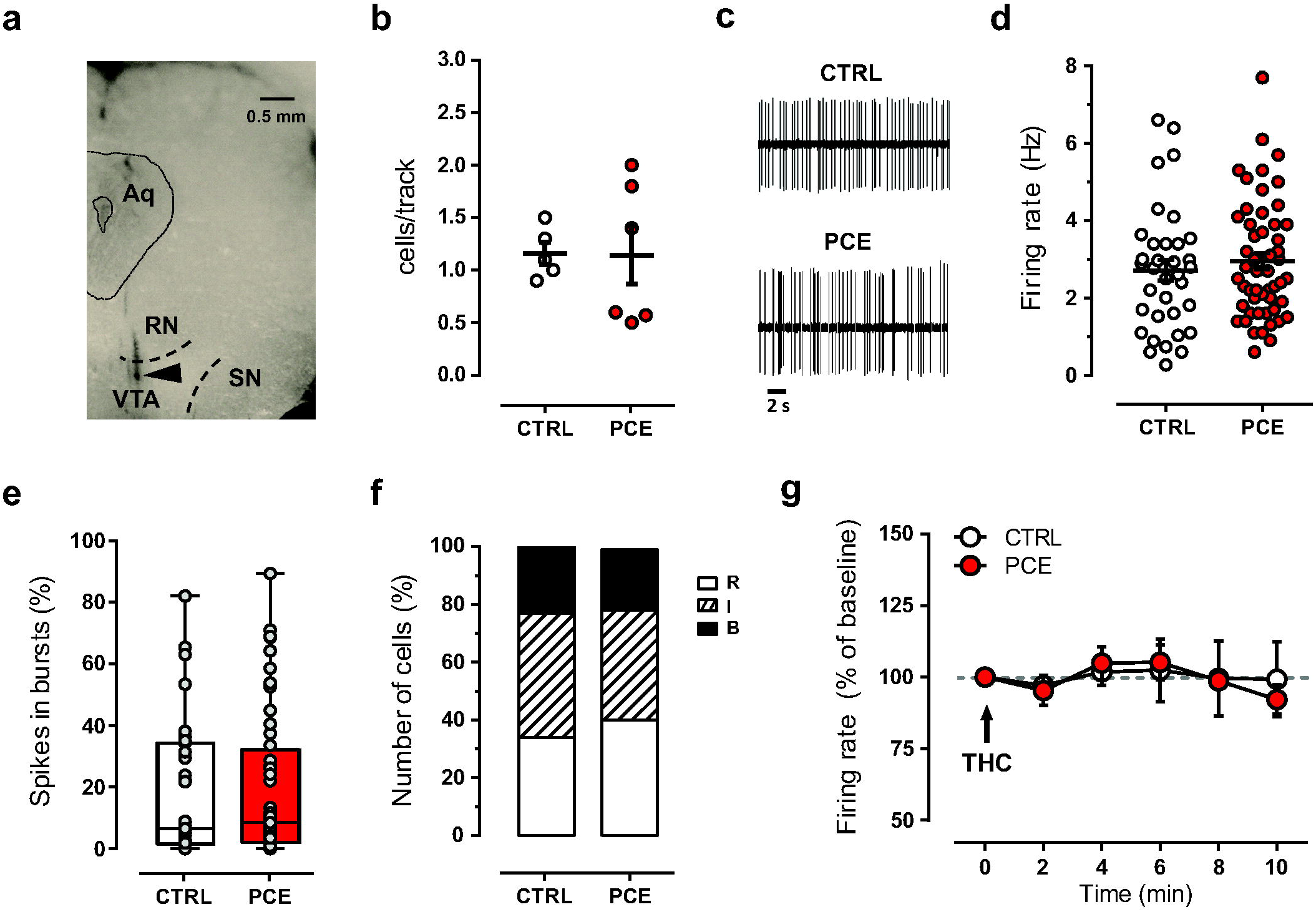
PCE effect on electrophysiological properties of putative dopamine neurons recorded in vivo from female preadolescent rats. (**a**) Coronal midbrain section from a preadolescent female rat showing the recording site (black triangle) in the ventral tegmental area. Abbreviations: Aq, aqueduct; RN, red nucleus; SN, substantia nigra, VTA, ventral tegmental area. (**b**) Scatter plot showing the average number of spontaneously active VTA dopamine neurons encountered per track. PCE (n_rats_=6) CTRL (n_rats_=5). (**c**) Representative traces of spontaneous firing activity of dopamine neurons from female offspring. (**d**) Spontaneous firing frequency of VTA dopamine cells from CTRL and PCE female rats. Data are represented as means s.e.m. with single values. (**e**) Percentage of spikes in burst displayed by dopamine neurons in female offspring. Data are represented as box and whisker plot with single values –min to max. (**f**) Stack bars represent the percentage of dopamine cells displaying different firing patterns: R=regular; I=irregular; B=bursty. (**g**) Time course of the effect of acute THC (0.5 mg/kg, i.v) on firing frequency of putative VTA dopamine neurons from PCE (n_cells_=5) and CTRL (n_cells_=5). Data represented as average ± s.e.m.

### 2.3 Intrinsic and synaptic properties of putative VTA dopamine neurons in PCE females

In male offspring, PCE imposes changes in both intrinsic and synaptic properties of putative dopamine neurons of the VTA by inducing a hyperexcitable phenotype ^34^. Ergo, we performed whole-cell patch clamp recordings from putative dopamine neurons of the lateral portion of the VTA in females. In acute brain slices, PCE did not impact VTA dopamine cell spontaneous firing rate (Figure 5a, b; unpaired t-test, t_29_=0.984, p=0.333) and their resting membrane potential (Figure 5c; unpaired t-test, t_31_=0.218, p=0.282). PCE did not change spike frequency in response to somatically injected currents (Figure 5d; linear regression, F_(1, 124)_= 1.757, P=0.187) and the latency to the first action potential (AP) elicited by the smallest current injected (Figure 5e; unpaired t-test, t_33_=0.765, p=0.449). No differences were also found in the voltage threshold of AP elicited by a depolarizing current between groups (Figure 5f; t_33_=0.577, p=0.567). We next examined synaptic properties of dopamine neurons by applying paired-pulse (50 ms interval) modulation protocol. PCE facilitated the paired pulse ratio (PPR) of AMPA-mediated excitatory postsynaptic currents (EPSCs) (Figure 6a; unpaired t-test, t_45_=2.444, p=0.018) without changing their coefficient of variation (CV) (Figure 6b; unpaired t-test, t_27_=1.35, p=0.188). No differences on current-voltage relationship of AMPA EPSCs (Figure 6c; linear regression, F_(1, 74)_= 0.001, p=0.971) and for AMPA/NMDA ratio were found between groups (Figure 6d; unpaired t-test, t_20_=0.901, p=0.378). Collectively, these findings indicate that PCE does not affect intrinsic properties and postsynaptic responsiveness to the glutamatergic transmission of VTA dopamine neurons of female offspring.

**Figure 5.**
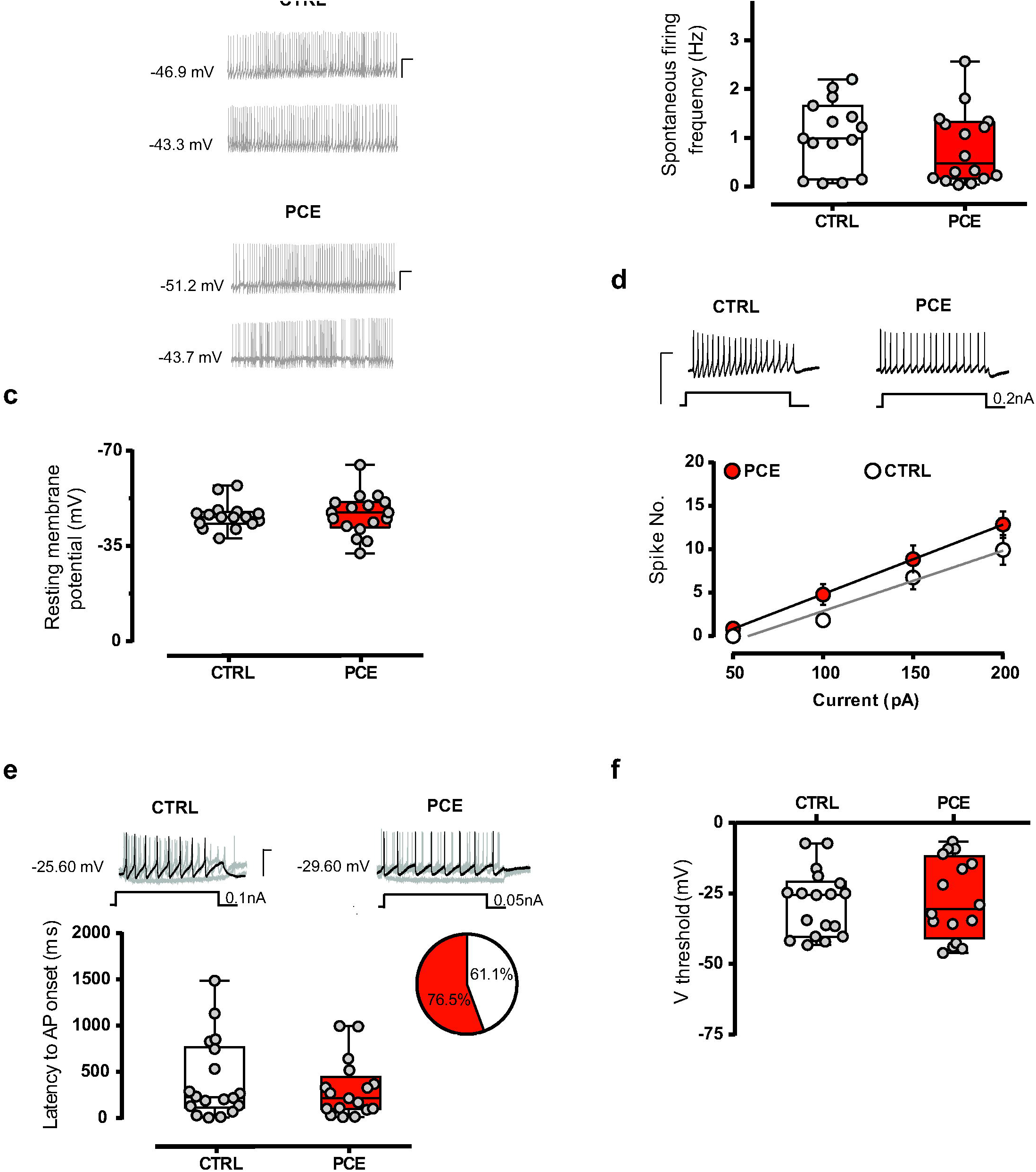
Intrinsic properties of *putative VTA dopamine neurons* are not affected by PCE in female preadolescent offspring. (**a**) Representative traces of spontaneous activity of dopamine neurons in acute VTA slices from CTRL and PCE offspring (*n*: 15 and 16 experiments from CTRL and PCE slices, respectively). Calibration bars, 100 ms, 20 mV. (**b,c**) PCE (*n*_*cells*_=16-17, *n*_*rats*_=11/group) does not affect the spontaneous firing frequency (**b**) and resting membrane potential (**c**) compared with CTRL cells (*n*_*cells*_=15-16, *n*_*rats*_=7/group). (**d**) Spike frequency in response to somatically injected current does not differ between CTRL (*n*_*cells*_= 18, *n*_*rats*_= 7/group) and PCE (*n*_*cells*_=17, *n*_*rats*_= 11/group) offspring. Insets show representative traces of evoked action potentials (APs) in response to maximum current injected. Calibration bar, 200 ms, 100 mV. Data are represented as average values per animal ± s.e.m. (**e**) Top: representative traces of evoked Aps in response to the minimum current injected. Calibration bar, 100 ms, 50 mV. Bottom: latency to the first AP of dopamine neurons elicited by the smallest current injected is not modify by PCE. Inset shows the proportion of cells eliciting APs at the smallest current (50 pA for PCE and 100 pA for CTRL group). (**f**) Threshold of AP elicited by depolarizing current does not vary between PCE (*n*_*cells*_=17, *n*_*rats*_=11/group) and CTRL (*n*_*cells*_=18, *n*_*rats*_=7/group) groups. Unless otherwise specified, all data are represented as box-and-whiskers plot with single values (min to max).

**Figure 6.**
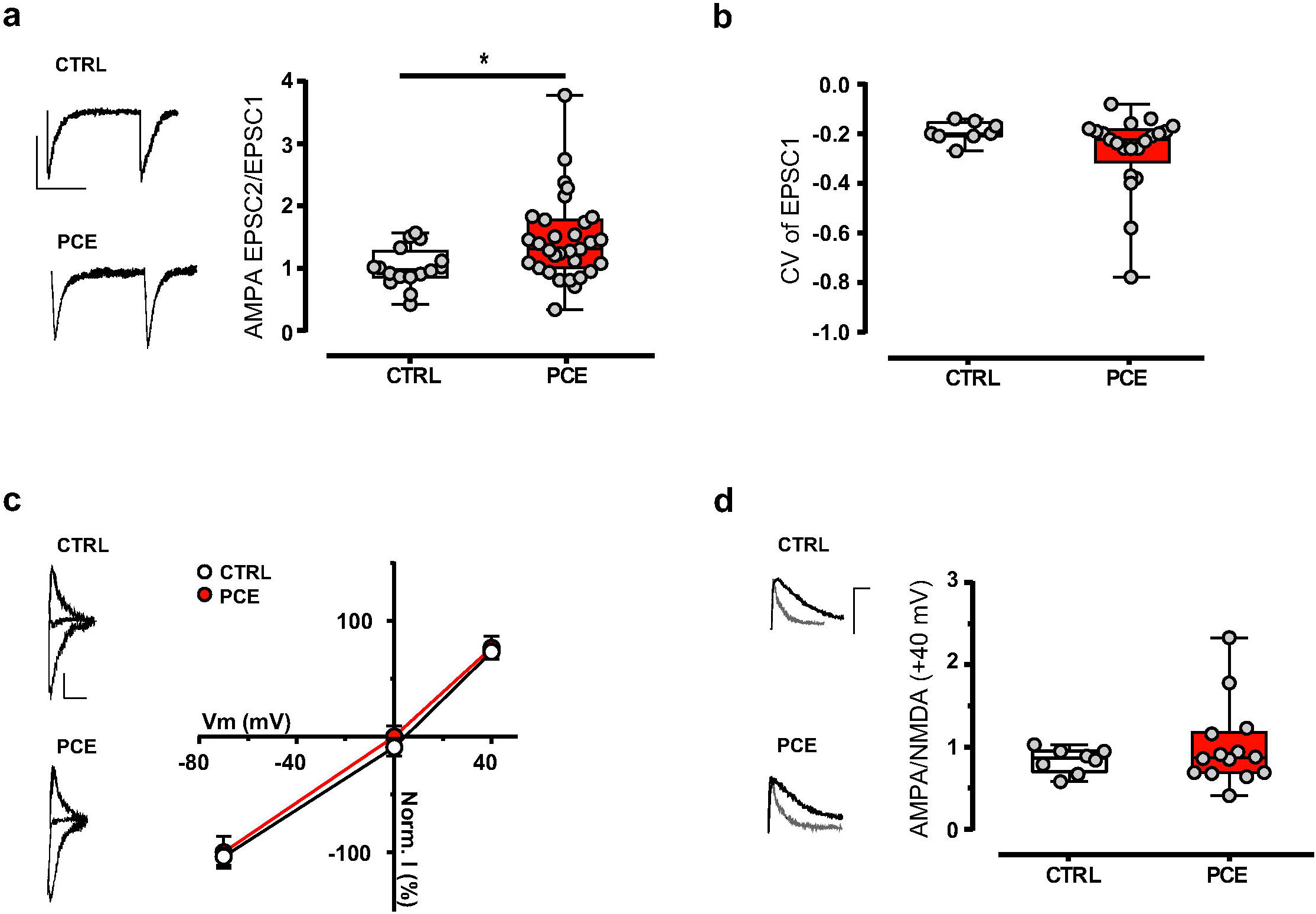
Excitatory synaptic properties of putative VTA dopamine neurons are not changed by PCE in female preadolescent offspring. **(a)** Dopamine cells from PCE (*n*_*cells*_: 31, *n*_*rats*_: 11/group) offspring exhibit an increased paired-pulse ratio (EPSC2/EPSC1) of AMPA EPSCs compared CTRL (*n*_*cells*_: 16, *n*_*rats*_: 7/group) group. Left-hand panel shows representative traces of paired AMPA EPSCs recorded from VTA putative dopamine neurons of CTRL and PCE offspring. Calibration bar: 25 ms, 50 pA. (**b**) PCE effects on 1/CV2 values from **a**. (**c**) Current-voltage relationship (I–V) curves of AMPA EPSCs recorded from dopamine neurons in PCE (*n*_*cells*_: 15, *n*_*rats*_: 12/group) and CTRL (*n*_*cells*_: 11, *n*_*rats*_: 8/group) offspring. Data are represented as mean ± s.e.m. AMPA EPSCs traces recorded at −70 mV, 0 mV and +40 mV from CTRL and PCE rats are shown on the left. Calibration bar: 10 ms, 25 pA. (**d**) The AMPA/NMDA ratio is not affected by PCE. Left panel shows representative traces of AMPA and NMDA EPSCs traces recorded from dopamine neurons held at +40mV in slices from PCE (*n*_*cells*_: 8, *n*_*rats*_: 11/group) and CTRL (*n*_*cells*_: 8, *n*_*rats*_: 7/group) offspring. Calibration bar: 10 ms, 50 pA. Unless otherwise specificated, all data are represented as box-and-whiskers plot with single values (min to max).

## 3. Discussion

The present study reveals a protection of pre-adolescent female progeny to prenatal THC exposure. Specifically, our findings show that not only PCE does not render females susceptible to acute THC ^34^, but we demonstrate that females show a resilient phenotype associated with normal mesolimbic dopamine (DA) system function. Specifically, PCE does not impact female offspring ability to cope with acute stress, to experience pleasure and to learn avoiding an unpleasant stimulus, thus broadening the spectrum of potentially disease-relevant behaviors examined that PCE could have been impacted.

Our findings support and extend previous animal studies where sex-specific differences in the effects of in utero exposure to THC, and more generally to cannabinoids, have been described ^32,35,38–41^. Remarkably, despite the importance of examining both sexes, this gap has only been partially recognized in research, but not yet fully addressed as analyzing only one sex or pooling the data from both sexes are still common practices. Nonetheless, evidence points to the impact of PCE being a function of sex despite the variables of the species, strain and age studied. On one hand, PCE’s detrimental effects on male sexual motivation, social interaction, spatial cognition and sensorimotor gating functions associated with alterations in the function of cortical and midbrain regions have been described^32,34,35,37,40,42^. On the other hand, in females, PCE affects spontaneous locomotion at adulthood ^38^ along with their motivation for food and morphine, an effect ascribed to changes in their mesolimbic DA activity ^41^. These contrasting findings arise from the many differences in experimental conditions among the studies but underline that sex-specific responses to PCE during fetal life do exist and are ascribed to differential effects on pathways and brain regions.

Our data, obtained during the prepuberal window of vulnerability, support previous studies examining different ages and showing that PCE does not affect female socioemotional behavior and coping strategies to acute stress, but decreases the reactivity of mesolimbic DA system as measured by the changes of extracellular dopamine levels in the nucleus accumbens shell (NAcS) in response to an acute challenge of THC. This latter, while in sharp contrast to our findings in male PCE counterparts ^34^, would be in agreement with the data showing in adult PCE female Wistar rats a reduction in the levels of 3,4-dihydroxyphenylacetic acid (DOPAC)/ DA ratio in both NAcS and VTA homogenates ^41^. DOPAC/DA ratio is a measure of DA turnover and has long been ascribed to mirror the activity of DA neurons in the VTA and the integrity of DA system function ^43^. However, when examined the DOPAC/DA ratio in our preadolescent PCE behaving Sprague Dawley female rats, we observed a two-fold increase in comparison with CTRL females at baseline, which subsided in response to an acute challenge with THC (data not shown). These discrepancies could be due to different strain of rats used, the age of the animals or the preparation for neurochemical analysis (behaving animals vs brain region homogenates). Additionally, our finding that firing frequency of VTA DA cells was not altered by PCE as indexed by our in vivo and ex vivo recordings, and that acute THC does not modify DA neuron spontaneous activity in vivo, support the notion that axonal DA release is not linearly scaling with somatic action potential firing ^44,45^. Although the latter is usually regarded as a proxy for dopamine signaling and its behavioral readout, many other factors regulate DA release in the target region, and at many stages ^46^, including DA production and vesicular loading, action potential propagation, regulation of dopamine reuptake, to name a few. To complicate this issue, sex differences in the mechanisms regulating DA release in the target regions are yet to be discovered. Finally, and importantly, sex dimorphisms in the mechanisms involved in neuronal intrinsic excitability and synaptic plasticity are seldom examined ^32,47^, especially before puberty; this is remarkable since sex hormones contribute to the differentiation of developmental trajectories and to dynamic changes of endocannabinoid signaling from adolescence to adulthood ^48^.

The observation of protection in females to the deleterious effects of PCE at pre-puberty supports the hypothesis that male sex is a risk factor for discrete neuropsychiatric disorders of developmental origin ^49–53^. One could speculate that because the growth rate of male fetuses is faster than females in the womb, the risk of undernutrition in males is increased. Notably, undernutrition is a contributing factor for the development of non-communicable diseases later in life ^54^. Fetal growth differences are usually associated with sex dimorphisms in placental expression of genes relevant for coping abilities to adverse in utero environment ^55,56^. However, there were no differences in weight gained by PCE offspring when compared to controls at pre-puberty in both females (data not shown) and males ^34^. Alternatively, one could argue that sex differences associated with global transcriptomic profiles and neuroprotective effects of glial cells in females ^57–59^ might protect them from the same environmental insult, i.e. THC. Noteworthy, the increased tightness of the blood-brain barrier in female neonatal rats ^59^ might protect them by reducing brain disposition of THC (or its metabolites), and the resulting interference with endocannabinoid system during fetal neurodevelopment. In particular, two xenobiotic transporters (i.e., Abcb1 and Abcg2) are involved in brain disposition of THC ^60^, and their expression shows sex dimorphisms in both the brain and the placenta ^61^. Indeed, female placentas express higher levels of Abcg2 mRNA than males ^61^, a transporter key in transplacental pharmacokinetics and fetal protection ^62^ that ensures proper function of fetoplacental unit throughout pregnancy ^63^. Since phytocannabinoids including THC inhibit the Abcg2 ^64^, one could speculate that brain THC concentrations might be higher in males than females, thus limiting its teratogenic impact on female neurodevelopment.

Collectively, our findings show that outcome of PCE in the female mesolimbic dopamine system outcome differs from that seen in males, though it might not strictly be a protected version of male outcome. Further investigations are needed to uncover potential PCE effects on females that might be region- and circuit- specific and associated with a disease-relevant phenotype potentially mutually exclusive. Finally, our results highlight the importance of consistently examining the mechanisms underlying physiological and pathological states in both sexes to develop tailored and personalized therapies.

## 4. Materials and Methods

### 4.1 Subjects

All experimental procedures were carried out according to the European legislation EU Directive 2010/63 and were approved by the Animal Ethics Committees of the University of Cagliari and by Italian Ministry of Health (auth. n. 256/2020). We made all efforts to minimize pain and suffering and to reduce the number of animals used.

Primiparous female Sprague Dawley rats (Envigo) were used as mothers and single housed during pregnancy. Offspring were weaned at post natal day (PND) 21 and were housed in in a climate-controlled animal room (21 ± 1 °C; 60% humidity) under a normal 12-h light–dark cycle (lights on at 7:00 a.m.) with *ab libitum* access to water and food. Because we previously found that PCE is a risk factor for psychotic-like endophenotype only in male progenies ^34^, the present investigation aims at testing whether female offspring shows a different disease-relevant behavioral phenotype linked to alterations in mesolimbic dopamine system function. Thus, all the experiments were conducted in female rats during pre-adolescence (PND15–28). To control for litter effects, no more than two females were used from each litter for the same experiment. To minimize the total number of animals used for the study, all the additional female pups in each litter were used for other experiments.

### 4.2 Drugs and Treatments

*Δ*9-Tetrahydrocannabinol (THC) resin was purchased from THC PHARM GmbH (Frankfurt, Germany) and dissolved in ethanol at 20% final concentration. Then, THC was suspended in a vehicle (VEH) solution containing 1–2% Tween^®^ 80 and diluted with sterile saline (0.9% NaCl).

To model PCE, rat dams were administered subcutaneously (s.c.) with THC (2 mg kg^−1^, 2 ml per Kg) or VEH once per day from GD5 until GD20. This dose of THC was chosen because it does not elicit substantial behavioral responses or tolerance after repeated administration ^65^. Moreover, it fails to affect maternal or non-maternal behavior, or offspring litter size ^34^. Notably, this dose of THC is equivalent to the current estimates of moderate cannabis consumption in human since it is similar to THC content in mild joints (5%) ^66^.

### 4.3 Behavioral Tests

#### Locomotor Activity

Thigmotaxis and motor behaviour of female offspring were recorded for 40 min in a novel, transparent open activity cage (Omnitech Digiscan monitoring). Female offspring (PND 24–28) were placed in a novel, square open-field arena (42 cm × 42 cm) surrounded by four 40-cm high transparent Plexiglas walls and locomotor activity was measured for 40 min using an Omnitech Digiscan monitoring system (Columbus, OH, USA) as indicated ^34^. Each cage (42 × 42 × 40 cm) had two sets of 16 photocells located at right angles to each other, projecting horizontal infrared beams 2.5□cm apart and 2□cm above the cage floor. 15 min of VEH or THC (2.5 mg kg−1, s.c.) administration, rats were placed in the center part of the arena and the total distance traveled, and the time spent in periphery and in center part of the arena were assessed.

#### Wire-beam bridge test

Risk propensity was evaluated using a variant of the wire-beam bridge task that was modified for rats, as detailed ^34^. Briefly, the apparatus consists of two Plexiglas platforms (156-cm high) connected by a horizontal, flexible wire-beam (100-cm long) bridge. One of the platforms was surmounted by a Plexiglas wall (52-cm high) placed right above the edge of the platform (start position). After 15 min of VEH or THC (2.5 mg kg^−1^, s.c.) administration, female offspring (PND 24-28) was individually placed in the start position and the entire session (3 min duration) was video-recorded. Behavioral measures included latency (s) to cross the bridge and reach the other platform and number of stretch-attend postures (SAPs) exhibited by animals during the test.

#### Elevated plus maze

The test was performed as previously described ^34^, under dim (10 lux) light. Briefly, the apparatus was made of black Plexiglas with a dark blue floor and consisted of two opposing open arms (length of 40 cm, width of 9 cm) and two closed arms (wall height of 15 cm), which extended from a central square platform (9 × 9 cm), positioned 70 cm high from the ground. Animals were tested after an acclimation period of 2 days in the experimental room. At the testing day, female rats (PND 24-28) were individually placed on the central platform facing the open arm. The entire sessions were video-recorded for 5 min and later scored by blinded observers. Behavioral measures included: time spent and entries into each partition of the maze. An arm entry was counted when all the four paws were inside the arm.

#### Forced Swim Test

To evaluate their responsiveness to an acute inescapable stressor, female offspring were tested by using a modified Porsolt forced swim test (FST) ^67^, as previously indicated ^37^. Rats (PND 24-28) were individually placed into a transparent cylinder (50 × 20 × 20 cm) filled with 2 l of cold water for 10 mins. During the test, animals were video-recorded and later scored by blinded observers. Behavioral measures included: passive coping (measured as duration of time spent floating with the absence of any movement except those necessary to keep the nose above water) and active coping behaviors (time spent swimming and climbing/struggling).

#### Passive Avoidance

Passive avoidance (PA) was tested in a two-way shuttle box using an experimental protocol modified for rats ^68^. The Apparatus consisted of two equal size (35 x 35 cm) compartments: light (white and illuminated with 24V-10W bulb) and dark (black and dark) divided by manually operated sliding door at the floor level. The dark compartment was equipped with an 18-bar insulated shock grid connected to a shocker. On day one (training session), female rats (PND 24-28) were individually placed in the light compartment facing away from the door and let to explore the chamber for 5 mins. The latency to enter into the dark compartment was measured. When the animal completely entered into the dark compartment, the sliding door was closed and a 0.5 mA shock was delivered for 2 s. After 30 s, the animal was removed from the apparatus and returned to its home cage. Animals that did not enter into the dark compartment within 5 mins were excluded from the analysis. After 24 hours (retention session), each animal was placed into the light chamber and the latency to enter the dark compartment was recorded to a maximum of 5 min.

#### Social Interaction Test

Social interaction was tested as previously described ^69,70^. Female rats (PND 24-26) were individually placed into a neutral, unfamiliar Makrolon cage (20 x 35 cm) together with a weight- and age-matched female conspecific (born from a separate litter) for 10 min. During the test animals were video-recorded and later scored by blinded observers. Behavioral measures included: total duration and frequency of social exploration (i.e. sniffing approaches directed to any part of the body of the partner), social play fighting behavior (i.e. pouncing, pinning, boxing, wrestling, or chasing) and nonsocial behaviors (i.e. exploratory activities directed to environment, grooming or digging activities).

#### Sucrose Preference Test

Anhedonia was measured by using a modified protocol of Sucrose Preference Test (SPT) ^71^. Before the test, female offspring were habituated to the presence of two drinking bottles for 2 days after weaning. Then (PND 23), rats were individually housed in a home cage with 2 drinking bottles: one containing 1% sucrose solution and the other with tap water for 5 days. The location of the bottles werewas switched daily in order to reduce habituation bias. Experimental measures included: Sucrose preference as percent (%) and averaged over the 5 days of testing. Sucrose Preference was calculated as a percentage of the volume of sucrose intake over the total volume of fluid intake.

### 4.4 Cerebral microdialysis

For cerebral microdialysis experiments, female rats were anesthetized with Equithesin and stereotaxically implanted with in-house constructed vertical microdialysis probes (AN 69-HF membrane, Hospal-Dasco; cut-off 40,000 Dalton, 3-mm dialyzing membrane length) in the nucleus Accumbens Shell (from bregma: anterior–posterior: + 1.5; lateral: ± 0.7; ventral: −7.0). The day after probe implantation, artificial cerebrospinal fluid solution (ACSF; 147 mM NaCl, 4 mM KCl, 1.5 mM CaCl2, 1 mM MgCl2, pH 6–6.5) was pumped through the dialysis probes at a constant rate of 1.1 μl min^−1^ via a CMA/100 microinjection pump (Carnegie Medicine, Stockholm, Sweden) in in freely moving animals. Samples were collected every 20 min and analyzed for dopamine content by high-performance liquid chromatography with electrochemical detection, as previously described ^72^. When a stable baseline was obtained (three consecutive samples with a variance not exceeding 15%), THC (2.5 mg per kg, 2 ml per kg) was intraperitoneally (i.p.) administered, and sample collection continued for 2 h. On completion of the experiments, rats were killed via an Equithesin overdose, the brains were removed and sectioned using a cryostat (Leica CM3050 S) into 40-μm thick coronal slices to verify the anatomical locations of dialysis probes.

### 4.5 In vivo single unit electrophysiological recordings

PND 25-29 female offspring were anaesthetized with chloral hydrate (400 mg/kg i.p.) and were cannulated in their femoral vein for intravenous administration of drugs. Animals were placed in the stereotaxic apparatus with their body temperature maintained at 37±1°C by a heating pad. Single unit activity was extracellularly recorded with glass micropipettes filled with 2% Pontamine sky blue dissolved in 0.5 M sodium acetate from lateral posterior VTA (anterior-posterior −4.5 to −5.2 mm from bregma, medial-lateral 0.4-

0.6 mm, ventral −6.5 to −7.5 mm from surface). Putative dopamine neurons were isolated and identified according to well established electrophysiological criteria such as their firing rate (<10 Hz), and duration of action potential (>2.5 ms). For each neuron, firing rate (calculated as the total number of spikes occurring over time) and bursting activity (defined as the occurrence of two spikes at inter-spike interval <80 ms and terminated when the inter-spike interval >160 ms) have been analyzed. In a subset of experiments, THC 0.5 mg/kg/ml was administered intravenously. At the end of recording sessions, a 15 mA current has been passed for 15 min through the micropipette in order to mark the position of the electrode within the recording site ^37^.

### 4.6 Ex vivo electrophysiological recordings

The preparation of posterior VTA slices was performed as previously described ^34^. Briefly, a block of tissue containing the midbrain was obtained from female offspring deeply anesthetized with isoflurane and the tissue sliced in the horizontal plane (300 µm) with a vibratome (Leica) in ice-cold low-Ca^2+^ solution containing the following (in mM): 126 NaCl, 1.6 KCl, 1.2 NaH_2_PO_4_, 1.2 MgCl_2_, 0.625 CaCl_2_, 18 NaHCO_3_ and 11 glucose. Slices were transferred to a holding chamber with ACSF (36-37 °C) saturated with 95% O_2_ and 5% CO_2_ containing the following (in mM): 126 NaCl, 1.6 KCl, 1.2 NaH_2_PO_4_, 1.2 MgCl_2_, 2.4 CaCl_2_, 18 NaHCO_3_ and 11 glucose. Slices were allowed to recover for at least 1 h before being placed, as hemislices, in the recording chamber and superfused with ACSF (36–37 °C) saturated with 95% O_2_ and 5% CO_2_. Cells were visualized using an upright microscope with infrared illumination (Axioskop FS 2 plus, Zeiss), and whole-cell patch-clamp recordings were made using an Axopatch 200B amplifier (Molecular Devices). Recordings were carried out in the lateral portion of the posterior VTA.

Voltage-clamp recordings of evoked EPSCs were made with electrodes filled with a solution containing the following (in mM): 117 cesium methanesulfonic acid, 20 HEPES, 0.4 EGTA, 2.8 NaCl, 5 TEA-Cl, 0.1 mM spermine, 2.5 Mg_2_ ATP and 0.25 Mg_2_ GTP, pH 7.2–7.4, 275–285 mOsm. Picrotoxin (100 μM) was added to the ACSF to block GABA_A_-receptor-mediated IPSCs. Series and input resistance were monitored continuously online with a 5-mV depolarizing step (25 ms). Current-clamp recordings were made with electrodes filled with a solution containing the following (in mM): 144 KCl, 10 HEPES buffer, 3.45 BAPTA, 1 CaCl_2_, 2.5 Mg_2_ATP and 0.25 Mg_2_GTP, pH 7.2–7.4, 275–285 mOsm. As previously described^1,11^, this solution had no effect on the holding current of the dopamine cells. Current-clamp experiments were performed in the absence of any pharmacological blocker, that is, in regular ACSF. Experiments were begun only after series resistance had stabilized (typically 10–30 MΩ), which was monitored by a hyperpolarizing step of −4 mV at each sweep every 10 s. Data were excluded when the resistance changed >20%. Data were filtered at 2 kHz, digitized at 10 kHz and collected online with acquisition software (pClamp 10.2, Molecular Devices).

Dopamine neurons from the lateral portion of the posterior VTA were identified according to previously published criteria ^34^ as follows: cell morphology and anatomical location (that is, medial to the medial terminal nucleus of the accessory optic tract); slow pacemaker-like firing rate (<5 Hz); long action potential duration (>2 ms) and the presence of a large hyperpolarization-activated current (I_h_> 100 pA) ^73^, which was assayed immediately after break-in using 13 incremental 10-mV hyperpolarizing steps (250 ms) from a holding potential of −70 mV. A bipolar, stainless steel stimulating electrode (FHC) was placed ∼100–200 μm rostral to the recording electrode and was used to stimulate at a frequency of 0.1 Hz.

Paired-pulse ratio (PPR), with an interstimulus interval of 50 msec, was calculated as the ratio between the second and the first postsynaptic currents (EPSC2/EPSC1) and averaged over 5 min. NMDA EPSCs were evoked while holding cells at +40 mV. The AMPA EPSC was isolated after bath application of the NMDA antagonist D-2-amino-5-phosphonovaleric acid (D-AP5, 100 μM). The NMDA EPSC was obtained by digital subtraction of the AMPA EPSC from the dual (AMPA + NMDA-mediated) EPSC ^34^.

### 4.7 Statistical analysis

Statistical analysis was performed with GraphPad Prism 6 (San Diego CA, USA) software. Data from behavioral, microdialysis and in vivo electrophysiological experiments were analyzed using two-tailed unpaired t-test or two-way ANOVA (followed by Bonferroni’s multiple comparisons test) when appropriated. Electrophysiological data were analyzed using Student’s t-test or linear regression when appropriate. Sample size was computed based on power calculations; The analysis assumptions are power = 0.9 and alpha = 0.5. Statistical outliers were identified with the Grubb's test (α = 0.05) and excluded from the analysis. Significance level was set at p < 0.05.

## Abbreviations

THC: Δ9-tetrahydrocannabinol

## Author Contributions

Conceptualization, M.M., R.F. and J.F.C.; methodology, F.T., C.S., V.S., M.C., P.S.; formal analysis, M.M., F.T., C.S., M.C., V.S.; resources, M.M.; data curation, F.T., C.S., P.D.; writing—original draft preparation, M.M.; writing—review and editing, J.F.C., R.F., P.D.; super-vision, M.M.; project administration, M.M.; funding acquisition, M.M., and J.F.C. All authors have read and agreed to the published version of the manuscript.

## Data Availability Statement

The data presented in this study are available on request from the corresponding author.

## Acknowledgments

We thank M. Tuveri, S. Aramo, and B. Tuveri for their skillful assistance.

